# Integrative analysis reveals genomic and epigenomic signatures of super-enhancers and its constituents

**DOI:** 10.1101/105262

**Authors:** Aziz Khan, Xuegong Zhang

**Author notes:** **Author’s email address**.

## Abstract

**Background:** Super-enhancers are clusters of transcriptional enhancers densely occupied by the Mediators, transcription factors and chromatin regulators. They control the expression of cell identity genes and disease associated genes. Current studies demonstrated the possibility of multiple factors with important roles in super-enhancer formation; however, a systematic analysis to assess the relative importance of chromatin and sequence signatures of super-enhancers and their constituents remain unclear.

**Results:** Here, we integrated diverse types of genomic and epigenomic datasets to identify key signatures of super-enhancers and their constituents and to investigate their relative importance. Through computational modelling, we found that Cdk8, Cdk9 and Smad3 as new key features of super-enhancers along with many known features such as H3K27ac. Comprehensive analysis of these features in embryonic stem cells and pro-B cells revealed their role in the super-enhancer formation and cellular identity. We also observed that super-enhancers are significantly GC-rich in contrast with typical enhancers. Further, we observed significant correlation among many cofactors at the constituents of super-enhancers.

**Conclusions:** Our analysis and ranking of super-enhancer signatures can serve as a resource to further characterize and understand the formation of super-enhancers. Our observations are consistent with a cooperative or synergistic model underlying the interaction of super-enhancers and their constituents with numerous factors.

## Background

Enhancers are *cis*-regulatory regions in the DNA that not only augment the transcription of associated genes but also play a key role in cell-type-specific gene expression [1,2]. A myriad of transcription factors (TFs) bind to enhancers and regulate gene expression by recruiting coactivators and RNA polymerase II (RNA Pol II) to target genes [3–7]. A typical mammalian cell is estimated to have thousands of active enhancers, a number which rises to roughly one million in the human genome [1,8]. It has been more than three decades since the first enhancer was discovered [9], but our understanding of the mechanisms by which enhancers regulate gene expression is still limited. However, development of methods such as chromatin immunoprecipitation followed by sequencing (ChIP-seq) and DNase I hypersensitivity followed by sequencing (DNase-seq), have helped to discover and characterize enhancers at genome scale. Many factors have been associated with enhancer activity, including mono methylation of histone H3 at lysine 4 (H3K4me1), acetylation of histone H3 at lysine 27 (H3K27ac), binding of the coactivator proteins p300 and CBP, and DNase I hypersensitivity [8,10,11]. By exploiting these factors and other genomic features, many computational approaches have been developed to predict enhancers genome-wide [4,12].

Mediator, a transcriptional coactivator, forms a complex with Cohesin to create cell-type-specific DNA loops and by facilitating enhancer-bound transcription factors, to recruit RNA Pol II to the promoters of target genes [13,14]. In embryonic stem cells (ESC), the pluripotency transcription factors Oct4, Sox2 and Nanog (OSN) co-bound regions have shown robust enhancer activity (25 out of 25) [15]. By using ChIP-seq data for Oct4, Sox2 and Nanog in ESC, 10,227 co-bound regions have been identified and classified into super-enhancers (SEs) or typical enhancers (TEs) by using ChIP-seq signal for Mediator subunit Med1 [16]. Super-enhancers form clusters of active enhancers, are cell-type specific, associated with key cell identity genes, and linked to many biological processes which define the cell identity [16]. These super-enhancers are densely loaded with the Mediator complex, master transcription factors and chromatin regulators [16–19]. Many disease- and trait-associated single nucleotide polymorphisms (SNPs) have been found in these regions [18]. Superenhancers differ from typical enhancers in terms of size, ChIP-seq density of various cofactors, DNA motif content, DNA methylation level, enhancer RNA (eRNA) abundance, ability to activate transcription and sensitivity to perturbation [16–18,20–22]. Further, studies have found superenhancers in multiple cancers and demonstrated their importance in cellular-identity and disease and emphasized their use as potential biomarkers [17,18,23–25]. Other parallel studies demonstrated nearly similar patterns by using different approaches and termed them ‘stretch enhancers’ [26] and ‘enhancer clusters’ [27].

Since the discovery of super-enhancers, the research community used ChIP-seq data for different factors to differentiate super-enhancers from typical enhancers in different cell-types. ChIP-seq data for Med1 optimally differentiated super-enhancers and typical enhancers by comparing it with enhancer marks, including H3K27ac, H3K4me1 and DNase I hypersensitivity [16]. BRD4, a member of the BET protein family, was also used to distinguish super-enhancers from typical enhancers as it is highly correlated with MED1 [17]. H3K27ac was extensively used to create a catalogue of super-enhancers across 86 different human cell-types and tissues due to its availability [18]. Other studies used the coactivator protein P300 to define super-enhancers [28,29]. However, the knowledge about these factors’ ability to define a set of super-enhancers in a particular cell-type and their relative and combinatorial importance remains limited. Master transcription factors which might form the superenhancer domains are largely unknown for most of the cell-types. Current studies demonstrated the possibility of multiple cofactors with important roles in super-enhancer formation; however, a detailed analysis is needed that integrates various types of data to investigate the key signatures of superenhancers and their constituents (enhancers within a super-enhancer). In addition, the degree to which the sequence-specific features of constituents by itself explains the differences between superenhancers and typical enhancers remains unknown.

Herein, to identify key features of super-enhancers and to investigate their relative contribution to predict super-enhancers, we integrated diverse types of publicly available datasets, including ChIP-seq data for histone modifications, chromatin regulators and transcription factors, DNase I hypersensitive sites and genomic data. Through correlation analysis and computational modelling, we showed that Med1, Med12, H3K27ac, Brd4, Cdk8, Cdk9, p300 and Smad3 are highly correlated and are more informative features to define super-enhancers. We performed a genome-wide analysis of Cdk8, Cdk9 and Smad3 binding in mESC and pro-B cells to analyse and assess their relative importance in defining super-enhancers. By using ChIP-seq data, RNA-seq based gene expression data, Gene Ontology (GO) and motif analyses we found that these factors may optimally differentiate super-enhancers from typical enhancers. Our systematic features analysis and ranking can be used as a platform to define and understand the biology of super-enhancers in other cell-types.

## Results

### Chromatin regulators and transcription factors at super-enhancer constituents

Studies have shown that super-enhancers are occupied by various cofactors, chromatin regulators, histone modifications, RNA polymerase II and transcriptions factors [16,18]. An understanding of the occupancy of these factors at the constituents of super-enhancers and typical enhancers is lacking. We extensively analysed 32 publicly available ChIP-seq and DNase-seq datasets to unveil their association with the constituents of super-enhancers and typical enhancers in mouse embryonic stem cells (mESC). We found that most of these factors, which are enriched in super-enhancers, were also highly enriched in the constituents of super-enhancers relative to typical enhancers (Fig. 1a; Figure S1 in Additional file 1). It is understandable to see that Oct4, Sox2 and Nanog were nearly equally enriched across the constituents of super-enhancers and typical enhancers because these constituents are defined by intersecting OSN co-bound regions [16].

**Fig. 1:**
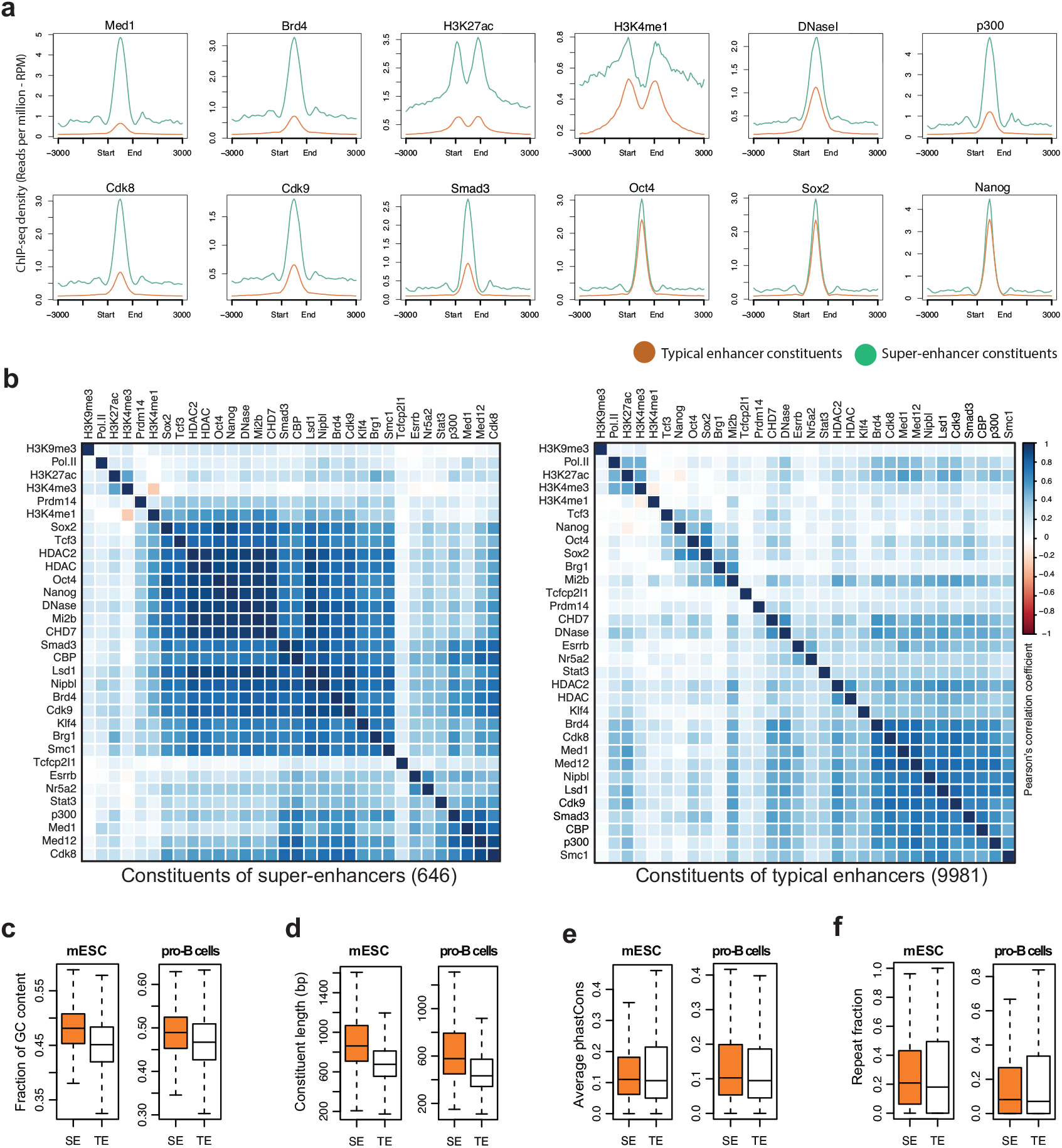
Chromatin regulators, transcription factors and sequence signatures of super-enhancer constituents. **(a)** Average ChIP-seq profile (RPM) of Med1, Brd4, H3K27ac, H3K4me1, DNaseI, p300, Cdk8, Cdk9, Smad3, Oct4, Sox2 and Nanog at the constituents of super-enhancers and typical enhancers, and their flanking 3kb regions **(b)** Correlation plot using Pearsons’ correlation coefficient with hierarchical clustering of normalized ChIP-seq signals (rpm/bp) of 32 factors at the constituents of super-enhancers (646) and typical enhancers (9981). **(c)** Box plot shows the fraction of GC-content, across the constituents of super-enhancers and typical enhancers in mESC and pro-B cells (p-value < 2.2e-16, Wilcoxon rank sum test). **(d)** Constituent enhancers size (bp) in mESC and pro-B cells (p-value < 2.2e-16, Wilcoxon rank sum test). **(e)** Conservation score (phastCons) in mESC (p-value = 0.6285, Wilcoxon rank sum test) and in pro-B cells (p-value < 1.7e-4, Wilcoxon rank sum test). **(f)** Repeat fraction in mESC (p-value = 0.0202, Wilcoxon rank sum test), pro-B cells (p-value = 0.8976, Wilcoxon rank sum test).

Through correlation analysis, we found that most of the factors were highly correlated at the constituents of super-enhancers compared to typical enhancers (Fig. 1b). Interestingly, features with similar lineage/functionality were clustered together. For example, histone modifications (H3K27ac and H3K4me3), Mediator complex subunits (Med1, Med12 and Cdk8), the pluripotency genes of ESC (Oct4, Sox2, Nanog) were clustered together. It was particularly interesting to observe that Smad3 clustered together with the co-activator protein p300/CBP. Previous studies have shown that p300/CBP interacts with Smad3 [30,31]. This suggests a possible combinatorial interplay among these cofactors at super-enhancers and that could be the reason that super-enhancers are more active and sensitive to perturbation as compared to typical enhancers.

### Sequence signatures of super-enhancers and their constituents

Super-enhancers differ from typical enhancers in terms of size, ChIP-seq density of various cofactors, TF content, ability to activate transcription, and sensitivity to perturbation [16–18]. But, to what extent constituents of super-enhancers differ from the constituents of typical enhancers in terms of sequence composition remains unknown. To gain insights into their biological functions, we sought to identify DNA sequence signatures of constituent enhancers. We tested GC-content, repeat fraction, size, and phastCons across the constituents of super-enhancers and typical enhancers in mouse ESC and pro-B cells.

Previous studies have shown that GC-rich regions have distinct features including frequent TF binding [32], active conformation [33] and nucleosome formation [34]. We found that constituents of super-enhancers are significantly more GC-rich than the constituents of typical enhancers (p-value < 2.2e-16, Wilcoxon rank sum test) (Fig. 1c). This suggests that GC content has an important role in super-enhancers formation and it can be a defining feature to distinguish them from typical enhancers.

Enhancers, larger than 3 kb have been shown to be cell-type-specific and are known as stretch enhancers [26]. A previous study showed that majority of super-enhancers do overlap with stretch enhancers [35]. We checked the size (bp) of constituents and found that constituents of superenhancers are significantly larger in size than the constituents of typical enhancers (p-value < 2.2e-16, Wilcoxon rank sum test) (Fig. 1d).

Enhancers are hardly conserved across mammalian genomes and evolved recently from ancestral DNA exaptation, rather than lineage-specific expansions of repeat elements [36]. We did not find any significant difference in conservation at constituents of super-enhancers and typical enhancers in mESC (p-value = 0.6285, Wilcoxon rank sum test), but in pro-B cells the conservation scores were slightly but significantly higher (p-value < 1.7e-4, Wilcoxon rank sum test) (Fig. 1e), which may suggest that super-enhancers are evolutionary conserved in differentiated cells. Similarly, there was no significant difference in repeat fraction at constituents of super-enhancers and typical enhancers in mESC (p-value = 0.0202, Wilcoxon rank sum test) or pro-B cells (p-value = 0.8976, Wilcoxon rank sum test) (Fig. 1f).

### Feature-ranking revealed previously known and new signatures of super-enhancers

With the increasing discovery of factors associated with super-enhancers, the determination of their relative importance in defining super-enhancers is important. Hence, we ranked chromatin and transcription factors to find a minimal optimal subset, which can be used to optimally distinguish super-enhancers from typical enhancers. We used a random-forest based approach Boruta [37] to assess the importance of each feature by ranking them based on their predictive importance (Fig. 2a) (Methods). We also used an out-of-bag approach to calculate the relative importance of each feature and achieved almost identical results (Figure S3 in Additional file 1).

**Fig. 2:**
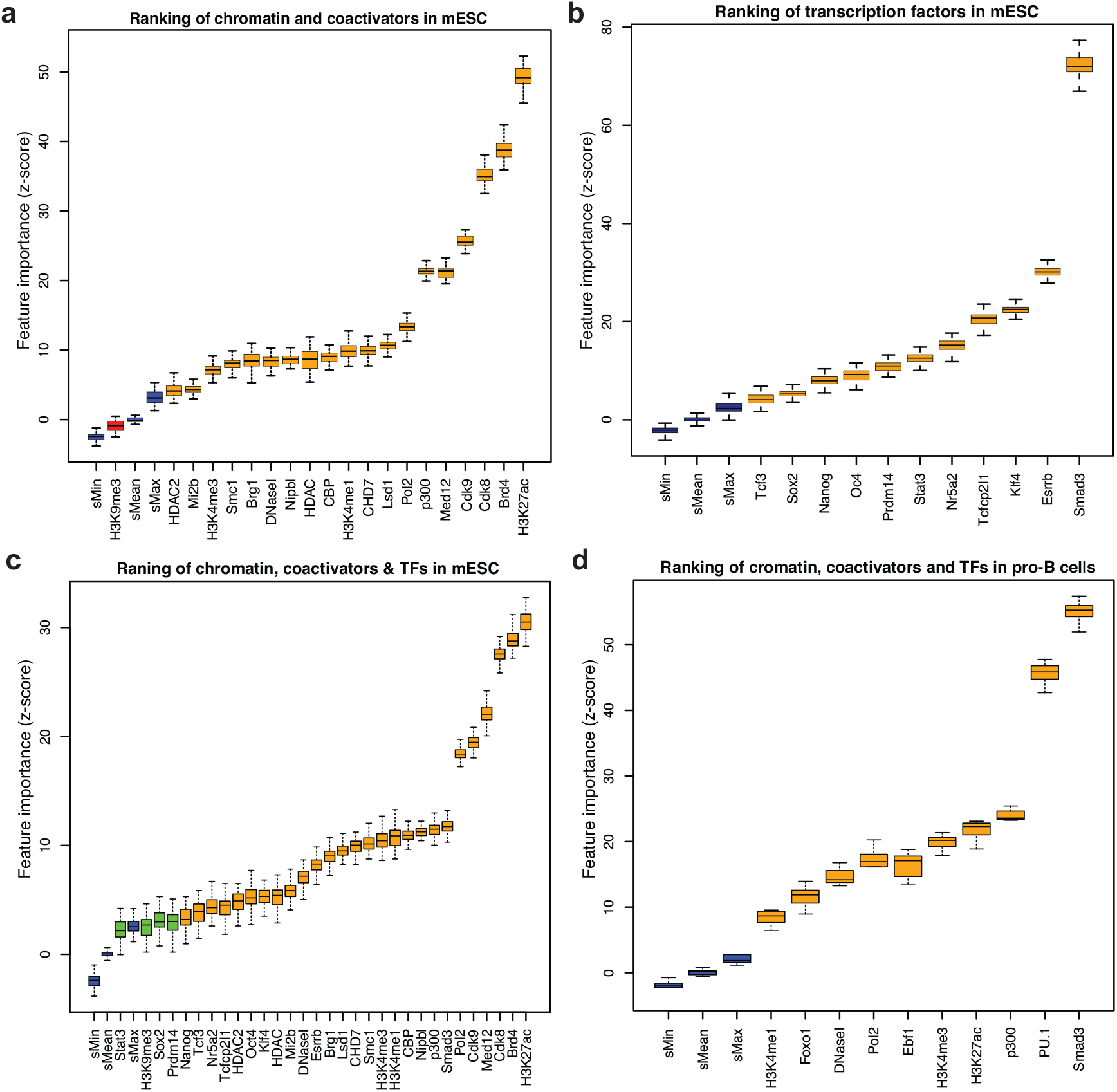
Ranking of features including chromatin and transcription factors in mESC and pro-B cells. **(a)** Box plot shows the importance of features, including histone modifications, chromatin regulators, coactivators, DNaseI and other features. The feature importance is calculated by using a Random Forest based algorithm, Boruta. The colors represents (Blue= shadow features; Red=negative features; Orange=Important features). **(b)** Box plot shows the feature importance for 11 transcription factors in mESC. **(c)** Box plot shows the importance of feature after combining the chromatin, coactivators and transcription factors features in mESC. **(d)** Box plot shows the importance features, including chromatin, coactivators and master transcription factors in pro-B cells.

After ranking chromatin features, we found Brd4, H3K27ac, Cdk8, Cdk9, Med12 and p300 as the six most important factors with Brd4 and H3K27ac as the top two most informative factors (Fig. 2a). It was particularly interesting to observe that Cdk8 and Cdk9 were ranked as the third and fourth most informative features, respectively. Cdk9, a subunit of the positive transcription elongation factor b (P-TEFb) has been found in enhancers and promoters of active genes along with the Mediator coactivator [17]. Cdk8, a subunit of Mediator complex, positively regulates precise steps in the assembly of transcriptional elongation, including the recruitment of P-TEFb and Brd4 [38].

Previous studies have shown that five ESC transcription factors (Sox2, Oct4, Nanog, Esrrb and Klf4) and other TFs (Smad3, Stat3, Tcf3, Nr5a2, Prdm14 and Tcfcp2l1) were enriched in superenhancers [16,18]. As these transcription factors are specific for ESC biology, we ranked them separately from other cofactors to find their relative importance in ESC. Interestingly, Smad3 turn to be the most informative among other transcription factors including Klf4 and Esrrb which were previously described as key defining features of super-enhancers [16] (Fig. 2b). It will be interesting to further understand the importance of Cdk8, Cdk9 and Smad3 in the formation of super-enhancers.

We also ranked the chromatin and transcription factors together, we found H3K27ac, Brd4, Cdk8, Cdk9 as most informative features, while Smad3 was ranked higher than p300 and also ESC specific master TFs (Fig. 2c). To test this in more differentiated cell-types, we have ranked several factors including H3K27ac, H3K4me1, H3K4me3, p300, Smad3, PU.1, Foxo1, Ebf1, Pol2 and DNaseI in pro-B cell. Interestingly, we found that Smad3 turn to be the most informative feature followed by PU.1, p300 and H3K27ac (Fig. 2d). This suggests that Smad3 might be more informative feature to distinguish super-enhancers in differentiated cells.

### Genome-wide profiles of Cdk8, Cdk9 and Smad3 at super-enhancers

Through the ranking of chromatin and transcription factors, we found that Cdk8, Cdk9 and Smad3 were important features along many known signatures of super-enhancers, including H3K27ac, Brd4, Med12 and p300 which are well characterized at super-enhancers [16–18,28]. However, the genome-wide profiles of Cdk8, Cdk9 and Smad3 are not well characterized at super-enhancers. Hence, we investigated the genome-wide profiles of Cdk8, Cdk9 and Smad3 at super-enhancers identified by using Med1 in mESC. We found that, ChIP-seq binding sites of Cdk8, Cdk9 and Smad3 were highly co-localized with Med1, Brd4, H3K27ac, p300 and DNaseI (Fig. 3a, b). Like Med1, the ChIP-seq density for Cdk8, Cdk9 and Smad3 is exceptionally higher at super-enhancers compared to typical enhancers (Fig. 3c). The ChIP-seq density of Med1, Cdk8, Cdk9 and Smad3 at super-enhancers is significantly higher compared to typical enhancers (p-value < 2.2e-16, Wilcoxon rank sum test) (Fig. 3d). Similarly, Med1, Cdk8, Cdk9 and Smad3 are also enriched at ESC super-enhancers identified using Med1 (Fig. 3f). The ChIP-seq binding for Med1 and master TFs, including Sox2, Oct4 and Nanog, is exceptionally higher and forms clusters at super-enhancers, which are associated with cell-type-specific genes [16]. We found that the ChIP-seq binding sites for Cdk8, Cdk9 and Smad3 also form clusters at super-enhancer regions and associated with cell-type-specific genes. For example, the super-enhancer (mSE_00038) is associated with ESC pluripotency gene Sox2 (Fig. 3e). In another example, the super-enhancer (mSE_00085) is associated with the ESC pluripotency gene Nanog and the super-enhancer (mSE_00084) is associated with Dppa3 (developmental pluripotency associated 3) gene, which plays a key role in cell division and maintenance of cell pluripotency (Figure S2A in Additional file 1).

**Fig. 3:**
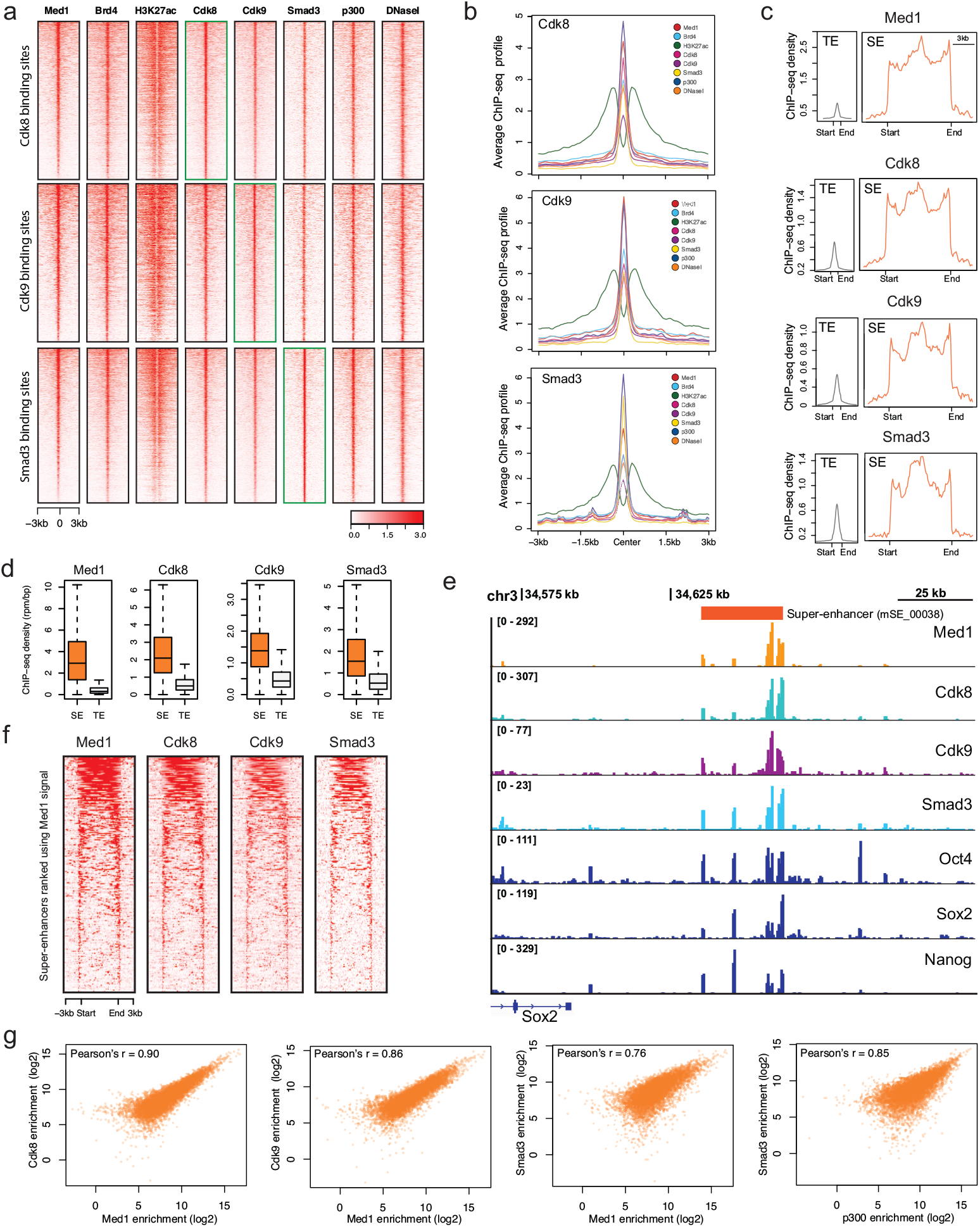
Genome-wide profiles of Cdk8, Cdk9 and Smad3 across super-enhancers and typical enhancers. **(a)** The heatmap shows the genome-wide ChIP-seq binding profile of Cdk8, Cdk9 and Smad3 across factors including Med1, Brd4, H3K27ac, Crdk8, Cdk9, Smad3, p300 and DNaseI. **(b)** The average read count profile of factors including Med1, Brd4, H3K27ac, Crdk8, Cdk9, Smad3, p300 and DNaseI across ChIP-seq sumits of Cdk8, Cdk9 and Smad3. **(c)** ChIP-seq density plots centred around super-enhancers and typical enhancers defined by Med1. Flanking regions are 3 kb. **(d)** Box plot shows the ChIP-seq density (rpm/bp) for Med1, Cdk8, Cdk9 and Smad3 in super-enhancers and typical enhancers defined by Med1. Box plot whiskers extend to 1.5x the interquartile range. **(e)** ChIP-seq binding profiles of Med1, Crdk8, Cdk9, Smad3, Oct4, Sox2 and Nanog at super-enhancer (mSE_00038) at the locus of Sox2 gene. **(f)** The heatmap of Med1, Crdk8, Cdk9 and Smad3 intensity at 231 mESC super-enhancers. **(g)** The left three scatter plot shows the Pearson’s correlation of Med1 with Cdk8, Cdk9 and Smad3 respectively at the OSN regions. The right most scatter plot shows the Pearson’s correlation between p300 and Smad3 at enhancer regions.

We calculated the Pearson’s correlation of Med1 with Cdk8, Cdk9 and Smad3 and found a high and significant correlation (p-value < 2.2e-16) (Fig. 3g). The correlation between Med1/Cdk8 (Pearson’s r = 0.90, p-value < 2.2e-16), Med1/Cdk9 (Pearson’s r = 0.86, p-value < 2.2e-16), and Med1/Smad3 (Pearson’s r = 0.76, p-value < 2.2e-16). Previous studies have used the ChIP-seq binding sites of the co-activator protein p300 to find enhancers [4,10]. We found that Smad3 is significantly correlated with p300 (Pearson’s r = 0.85, p-value < 2.2e-16) (Figure 4g).

**Fig. 4:**
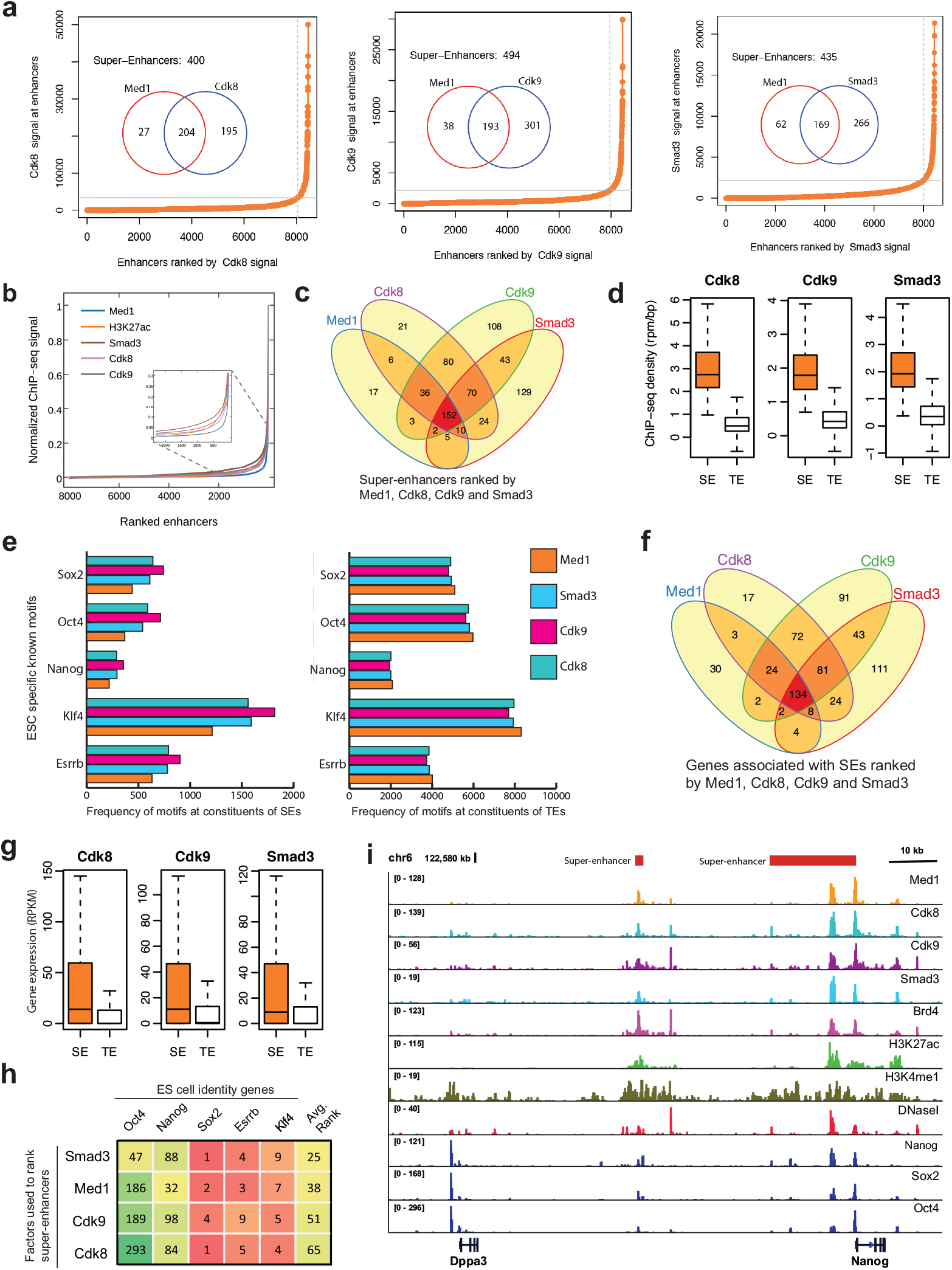
Super-enhancers identified by using Cdk8, Cdk9 and Smad3. **(a)** The Hockey-stick plot shows the cut-off used to separate super-enhancers from co-OSN (Oct4, Sox2, and Nanog) regions by using Cdk8, Cdk9 and Smad3. **(b)** The distribution of normalized ChIP-seq signal of Med1, H3K27ac, Cdk8, Cdk9 and Smad3 at mESC enhancers. For each factor, the values were normalized by dividing the ChIP-seq signal at each enhancer by the maximum signal. The rank of enhancer at each factor was measured independently. The figure zoomed at the cut-off so it can be visualized. **(c)** Venn diagram shows the number of super-enhancers overlapped, ranked using Med1, Cdk8, Cdk9 and Smad3. **(d)** Boxplot shows the ChIP-seq density (rpm/bp) in super-enhancers and typical enhancers defined by Cdk8, Cdk9 and Smad3 (p-value < 2.2e-16, Wilcoxon rank sum test). **(e)** Bar-plot shows the frequency of motifs (Oct4, Sox2, Nanog, Klf4 and Essrb) found at the constituents of super-enhancers and typical enhancers defined by Med1, Cdk8, Cdk9 and Smad3. **(f)** Venn diagram of genes associated with super-enhancers identified using Med1, Cdk8, Cdk9 and Smad3. The genes associated to Med1 super-enhancers are downloaded from dbSUPER. **(g)** Boxplot shows the gene expression (RPKM) in super-enhancers and typical enhancers defined by Cdk8, Cdk9 and Smad3. **(h)** Rank of factors based on the rank of super-enhancers associated with of ESC identity genes, including Sox2, Oct4, Nanog, Esrrb and Klf4. The table is sorted based on the average rank. **(i)** ChIP-seq binding profiles of different factors at the typical enhancer and super-enhancers at Dppa3 and Nanog gene locus in mESC.

### Identification and characterization of super-enhancers by using Cdk8, Cdk9 and Smad3 in mESC

Since we found that Cdk8, Cdk9 and Smad3 are highly correlated with Med1 and co-occupy superenhancers genome-wide (Fig. 3), we investigated the importance of Cdk8, Cdk9 and Smad3 in superenhancer formation and also compared them with super-enhancers identified by Med1. We used ChIP-seq data and RNA-seq data to identify and characterize super-enhancers by using Cdk8, Cdk9 and Smad3 in mESC. We found 400, 494 and 435 super-enhancers by using Cdk8, Cdk9 and Smad3, respectively (Fig. 4a). A list of all the super-enhancers and typical enhancers can be found in (Additional file 2). Furthermore, Cdk8, Cdk9 and Smad3 successfully identified 88%, 84% and 73% of the Med1 super-enhancers, respectively (Fig. 4a). After Med1, we can see more clear distinction of super-enhancers and typical enhancers by using Cdk8 as compared with H3K27ac (Figure 5b). The majority of super-enhancers identified using Cdk8, Cdk9 and Smad3 do overlap with super-enhancers identified using Med1, which composes 66% of the super-enhancers identified using Med1 (Fig. 4c).

**Fig. 5:**
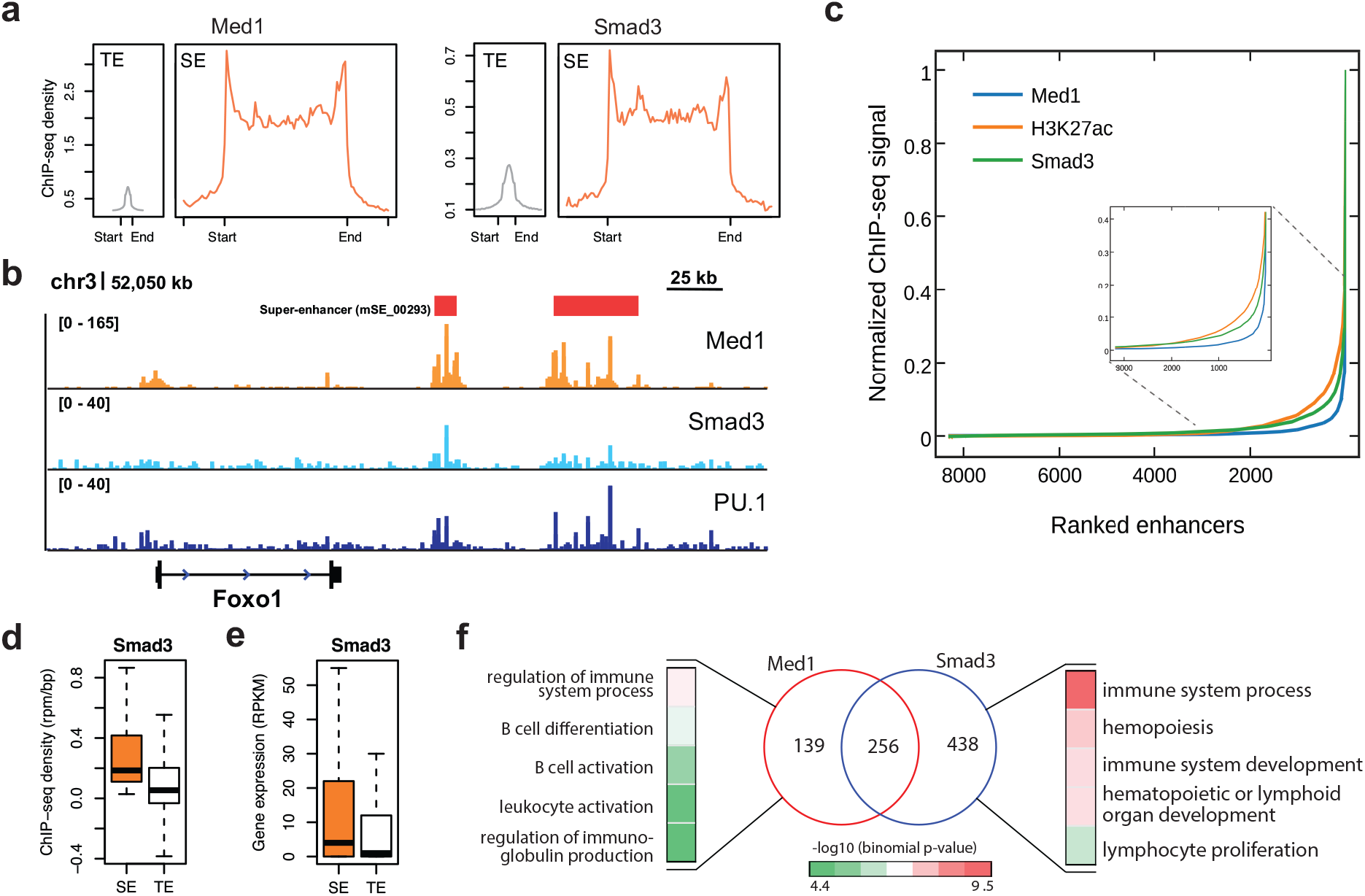
Comparison of super-enhancers ranked using Med1 and Smad3 in pro-B cells. **(a)** Average ChIP-seq density of Med1 and Smad3 across 13,814 typical enhancers and 395 super-enhancers identified using Med1. The flanking region is 3kb. **(b)** ChIP-seq binding profiles for Med1, Smad3 and PU.1 at the locus of Foxo1 gene. The super-enhancer (mSE_00293) is associated with Foxo1 gene. **(c)** The distribution of normalized ChIP-seq signal of Med1, H3K27ac and Smad3 at pro-B enhancers. For each factor, the values were normalized by dividing the ChIP-seq signal at each enhancer by the maximum signal. The rank of enhancer at each factor was measured independently. The figure zoomed at the cut-off so it can be visualized. **(d)** Box-plot shows the Smad3 ChIP-seq density (rpm/bp) at super-enhancers and typical enhancers regions defined using Smad3 in pro-B cells. **(e)** Box plot shows the gene expression (RPKM) for the genes associated with superenhancers and typical enhancers defined using Smad3 in pro-B cells. **(f)** The Venn diagrams show the overlap of super-enhancers identified using both Med1 and Smad3 in pro-B cells. Super-enhancer regions identified using Med1 in pro-B were obtained from [16]. Gene Ontology terms (Biological Process) for super-enhancers identified by Med1 only or Smad3 in pro-B cells.

The ChIP-seq density at super-enhancers identified using Cdk8, Cdk9 and Smad3 is significantly higher compared with typical enhancers (p-value < 2.2e-16, Wilcoxon rank sum test) (Fig. 4d).

In ESC, the DNA motifs Klf4 and Esrrb were particularly enriched at the constituents of superenhancers, compared to typical enhancers [16]. Hence, we tested the frequency of these two motifs at the constituents of super-enhancers and typical enhancers; we defined using Cdk8, Cdk9 and Smad3. We found that the frequency of binding motifs Klf4 and Esrrb is significantly higher at constituents of super-enhancers than typical enhancers (p-value < 2.2e-16, Wilcoxon rank sum test) (Figure S2G in Additional file 1). When we compared the frequency of known ESC specific motifs (Oct4, Sox2, Nanog, Esrrb and Klf4) at the constituents of super-enhancers and typical enhancers defined by Med1, Cdk8, Cdk9 and Smad3. We found a higher frequency of these motifs at superenhancers defined by Cdk9, Cdk8 and Smad3 as compared with Med1 (Fig. 4e). The frequency of these motifs was also slightly higher at the typical enhancers identified by Med1 as compared with Cdk8, Cdk9 and Smad3.

The genes associated with super-enhancers are highly expressed, compared to genes associated with typical enhancers [16–18]. To test this we associated genes with the super-enhancers and typical enhancers as described in [16,18]. We found that 65% of the Med1 super-enhancers associated genes were also associated with super-enhancers identified by Cdk8, Cdk9 and Smad3 (Fig. 4f). These genes associated with super-enhancers were significantly highly expressed, compared to genes associated with typical enhancers (p-value < 2.2e-16, Wilcoxon rank sum test) (Fig. 4g).

Super-enhancers known to be highly enriched for cell-type-specific master regulators and these regulators should have a higher rank. Hence, we checked the rank of super-enhancers, associated with the key cell identity genes, including Oct4, Sox2, Nanog, Esrrb and Klf4 in ESC, and ranked the factors based on the average rank of the super-enhancers that are associated with these genes (Fig. 4h). These genes were selected due to their important roles in the pluripotency and reprogramming of ESC biology [39–41]. The rankings for Med1 super-enhancers were downloaded from [18]. We found that Smad3 achieved a highest rank followed by Med1, Cdk9 and Cdk8. The Smad3 achieved a higher rank for Oct4, compared with other genes. This might be due to the fact that Smad3 co-bonded with the master transcription factor Oct4 genome-wide in ESC [42]. We found similar ChIP-seq patterns for factors including Med1, H3K27ac, Brd4, Cdk8, Cdk9 and Smad3 at super-enhancers regions defined by all three factors (Cdk8, Cd9 and Smad3) and are associated with cell-type-specific genes, including Nanog and Dppa3 (Fig. 4i). Like Med1, the genes associated with super-enhancers ranked by Cdk8, Cdk9 and Smad3 were enriched with cell-type-specific GO terms, supporting the notion that super-enhancers regulate cellular identity genes (Figure S4a in Additional file 1). Taken together, our results indicate the role of Cdk8, Cdk9 and Smad3 in defining and formation of superenhancers.

### Identification and characterization of super-enhancers by using Smad3 in pro-B cells

It was particularly interesting to see Smad3 ranked the most informative among transcription factors, including Oct4, Sox2, Nanog, Esrrb, Klf4, Tcfcp2l1, Prdm14, Nr5a2, Stat3 and Tcf3 in mESC (Fig. 2b). In ESC, the highly ranked super-enhancers identified using Smad3 were associated with ES cell-identity genes, including Oct4, Sox2, Nanog, Klf4 and Esrrb compared to Med1 super-enhancers (Fig. 4h). A previous study showed that Smad3 co-occupy sites with the master transcription factors (Oct4 in mES Myod1 in myotubes, and PU.1 in pro-B cells) genome-wide [42]. We also observed Smad3 turn to be the highly ranked feature when we ranked Smad3 and several other factors in pro-B cells (Fig. 2d).

Hence, we argued that Smad3 could be used to define super-enhancers where Med1 is not available. We already demonstrated above that Smad3 could be used to define super-enhancers in ESC. To test this in more differentiated cells, we identified and characterized super-enhancers in pro-B cells using Smad3. We compared the super-enhancers identified by using Smad3 with previously identified super-enhancers defined by using Med1 in pro-B cells [16]. The ChIP-seq density of Smad3 super-enhancers is exceptionally higher compared to typical enhancers (Fig. 5a). Smad3 have strong binding along with Med1 and PU.1 at a super-enhancer(mSE_00293) which is associated with Foxo1 gene (Fig. 5b).

By using Smad3, we identified 694 super-enhancers among which 65% were identified by Med1 (Figure S2f in Additional file 1). With Smad3 we can see a clearer distinction of super-enhancers and typical enhancers as compared with H3K27ac (Fig. 5c). The ChIP-seq density at super-enhancers identified using Smad3 is significantly higher, compared to typical enhancers (p-value < 2.2e-16, Wilcoxon rank sum test) (Fig. 5d). The genes associated with Smad3 super-enhancers are significantly expressed, compared with typical enhancers (p-value < 2.2e-16, Wilcoxon rank sum test) (Fig. 5e). The GO terms for super-enhancers ranked by Smad3 are highly enriched and cell-type-specific, compared to Med1 (Figure S4b in Additional file 1).

To test the functional importance of super-enhancers identified only by Smad3 or by Med1, we performed GO analysis on genes associated with this subset of super-enhancers. Interestingly, the super-enhancers identified by Smad3 but not by Med1 turn to be highly enriched for cell-type-specific GO terms such as immune cell development and immune system development. While superenhancers identified by Med1 but not by Smad3 lowly enriched for cell-type-specific GO terms (Fig. 5f). These results, taken together with previous studies, demonstrate the importance of Smad3 in super-enhancer formation and Smad3 could be used to define super-enhancers.

## Discussion

Super-enhancers regulate expression of key genes that are critical for cellular identity, thus, alterations at these regions can lead to several disorders. Hence, exploring these cis-regulatory elements and their features to uncover the molecular mechanisms that orchestrates the cell-type-specific gene regulation in development and disease.

In this study, we first presented a systematic approach to rank and access the importance of different features of super-enhancers. We investigated different features including histone modifications, chromatin regulators, transcription factors, DNA hypersensitive sites and DNA sequence motifs in mESC. We also analysed sequence-specific features including GC content, conversation score and repeat fraction in mESC and pro-B cells. We found new features including Cdk8, Cdk9, Smad3 and GC content as key features of super-enhancers along with many known features, which make super-enhancers distinct from typical enhancers.

Among the chromatin features we found that Brd4, H3K27ac, Cdk8, Cdk9, Med12 and p300 were the top six features of super-enhancers. Previous studies showed the importance of these highly ranked features and their role in transcriptional regulation. Through our ranking of features, H3K27ac achieved a higher ranking, compared to other histone modifications. H3K27ac was found as a mark to separate active enhancers from poised enhancers [11], which suggests that super-enhancers might be the clusters of active enhancers. The Mediator sub-units Med1 and Med12 has been known as master coordinators of cell lineage and development [16,43]. We did not include Mediator sub-unit Med1 in our feature ranking because a Med1 signal was used to define super-enhancers [16]. Bromodomain-containing protein 4 (Brd4), a member of the BET protein family which functions as an epigenetic reader and transcriptional regulator that binds acetylated lysines in histones [44], was ranked as the second highest important feature. Brd4 has been associated with anti-pause enhancers (A-PEs) which regulate the RNA Polymerase II (Pol II) promoter-proximal pause release [45,46]. Brd4 regulates the positive transcription elongation factor b (P-TEFb) to allow Pol II phosphorylation and the subsequent elongation of target genes [46,47]. In ESC it specifically governs the transcriptional elongation by occupying super-enhancers and by recruiting Mediator and Cyclin dependent kinase 9 (Cdk9) to these super-enhancers [48]. Cdk9, a sub-unit of P-TEFb, has been found at enhancers and promoters of active genes along with the Mediator coactivator [17]. Cyclin-dependent kinase 8 (Cdk8), a subunit of Mediator complex, positively regulates precise steps in the assembly of transcriptional elongation, including the recruitment of P-TEFb and BRD4 [38]. A very recent study has demonstrated that Cdk8 regulates the key genes associated with super-enhancers in acute myeloid leukaemia (AML) cells [49]. We identified and characterized super-enhancers in mESC by using Cdk8 and Cdk9, to further validate their importance in super-enhancer formation and cell identity.

Among the transcriptions factors, we found that Smad3, Esrrb, Klf4, Tcfcp2l1, Nr5f2a and Stat3 were the top ranked features of super-enhancers. It was particularly interesting to see that Smad3 was ranked as the best feature among the transcription factors including Esrrb and Klf4. We know that Smad3 is a target of the TGF-β signaling pathway, and studies have shown that Smad3 is recruited to enhancers formed by master transcription factors [42]. We found significant correlation between Smad3 and coactivators p300/CBP at super-enhancers and previous studies have shown that p300/CBP interacts with Smad3 [30,31]. The evidence for the enrichment of Smad3 at superenhancers shows how the transforming growth factor beta (TGF-β) signaling pathway can converge on key genes that control ES cell identity. A very recent study validates our findings by showing that super-enhancers provide a platform for signalling pathways, including TGF-β to regulate genes that control cell identity during development and tumorigenesis [50]. To validate further, we identified and characterized super-enhancers using Smad3 in mESC and pro-B cells. Our analysis through the integration ChIP-seq and RNA-seq data suggested the importance of Smad3 in super-enhancer formation and cell identity.

By investigating sequence signatures, we found that the constituents of super-enhancers were significantly GC-rich. The GC-richness of a genomic region is associated with several distinctive features that can affect the *cis*-regulatory potential of a sequence [32,33]. GC-rich and AT-rich chromatin domains are marked by distinct patterns of histone modifications. GC-rich chromatin domains tend to occur in a more active conformation and histone deacetylase activity represses this propensity throughout the genome [33]. Also GC content and nucleosome occupancy are positively correlated [32] and GC-rich sequences promote nucleosome formation [34]. Transcription factors tend to bind GC-rich regions in the genome, regardless of the distance and orientation [32]. This suggests that there is a role for the GC content in the formation of super-enhancers, which control the cell-type-specific gene expression.

Enhancers function due to cooperative and synergistic interplay of different coactivators and transcription factors [51]. A recent study showed that multiple enhancer variants cooperatively contribute to altered expression of their gene targets [52]. It is not well understood whether constituents of super-enhancers work synergistically or additively. The constituents of superenhancers make frequent physical contacts with one another [53] and extensive cooperative binding of transcription factors have been found at super-enhancers [54]. A study in ESC demonstrated the functional importance of super-enhancer constituents [50]. Recent studies have suggested additive and functional hierarchy among the constituents of α-globin and Wap super-enhancer locus, respectively [55,56]. But, a very recent study argues that it is still need to be determined whether the constituents of a super-enhancer functions synergistically or additively [57]. Through computational modelling and correlation analysis, we noticed a combinatorial relationship between chromatin regulators and transcription factors at the constituents of super-enhancers. This advanced our current understanding of the determinants of super-enhancers and led us to hypothesize that these combinatorial patterns may be involved in mediating super-enhancers. Furthermore, the significant correlation of many cofactors at the constituents of super-enhancers suggests cooperative and synergistic interactions. These results, taken together with previous studies suggest a cooperative and synergistic interplay of between constituents of super-enhancers. More sophisticated experiments are needed to validate the functional importance of constituents within a super-enhancer, by utilizing techniques such as CRISPR-Cas9.

## Conclusions

We integrated diverse types of genomic and epigenomics datasets to identify new signatures of super-enhancers and their constituents. We investigated the relative importance of each feature in predicting super-enhancers. We found Cdk8, Cdk9 and Smad3 as new signatures, which can be used to define super-enhancers where Mediator or master transcription data is not available. Taken together with previous studies, our results are consistent with a cooperative or synergistic model underlying the interaction of super-enhancers and their constituents with numerous factors. Our systematic feature analysis and ranking can serve as a resource to further characterize and understand the formation and molecular mechanisms of super-enhancers.

## Methods

### Data description

We downloaded 32 publicly available ChIP-seq and DNase-seq datasets in mouse embryonic stem cells (mESC) from Gene Expression Ominibus (GEO). These include four histone modifications: H3K27ac, H3K4me1, H3K4me3 and H3K9me3; DNA hypersensitive site (DNaseI); RNA polymerase II (Pol II); transcriptional co-activating proteins (p300, CBP); P-TFEb subunit (Cdk9); sub-units of Mediator complex (Med1, Med12, Cdk8); other chromatin regulators (Brg1, Brd4, Chd7); Cohesin (Smc1, Nipbl); subunits of Lsd1-NuRD complex (Lsd1, Mi2b) and 11 transcription factors (Oct4, Sox2, Nanog, Esrrb, Klf4, Tcfcp2l1, Prdm14, Nr5a2, Smad3, Stat3 and Tcf3). A detailed list of all datasets used in this study is provided in (Table S1 in Additional file 1).

To identify super-enhancers and perform features ranking in pro-B cells, we used ChIP-seq data for Med1, PU.1, Foxo1, Smad3, Ebf1, p300, H3K27ac, H3K4me1, H3K4me3 and Pol2 and also DNase-seq (Table S2 in Additional file 1).

We also obtained processed RNA-seq based gene expression data (RPKM) from [58] and [16] for mESC and pro-B cells, respectively. We downloaded super-enhancer regions in mESC and pro-B cells identified using Med1 ChIP-seq occupancy from dbSUPER, which were identified using similar approach [59]. We also used other genomic features including GC content, conservation score (phastCons) and repeat fraction downloaded from the UCSC table browser [60].

### Data analysis and feature extraction

Initially, ChIP-seq reads were aligned to mouse genome-build mm9 using bowtie [61] (Version 0.12.9) with parameters (-k 1, -m 1, -n 2, -e 70, –best). We calculated read densities for 30 ChIP-seq datasets at the constituents of super-enhancers (646) and typical enhancers (9981) and normalized it as described in [16,62]. Briefly, for each constituent region, reads were extended by 200bp and the density of reads per base pair was calculated using bamToGFF (https://github.com/BradnerLab/pipeline). Next, these densities were normalized in units of reads per million mapped reads per base pair (rpm/bp) with background subtraction. We used the similar approach for DNase-seq data but without background subtraction.

We used MACS (Model-based Analysis of ChIP-Seq) [63] (Version 1.4.2) to perform the peak calling and to find ChIP-seq-enriched regions over background. We used a p-value (10^−9^) as the enrichment threshold. To generate wiggle files, we used MACS with parameter -w -S --space=50.

### Identification of constituent enhancers and super-enhancers

We used 10,227 co-bound regions of Oct4, Sox2 and Nanog in ESC and 13,814 regions of master transcription factor PU.1 in murine progenitor B (pro-B) cells as constituent enhancers, which were obtained from [16]. These constituent enhancers were then stitched together with in 12.5 kb and +/-2 kb away from TSS. These stitched regions were ranked based on the ChIP-seq signal by using the ROSE (Rank Ordering of Super-Enhancers) software to define super-enhancers [16]. The superenhancers data downloaded from dbSUPER was also generated using the same pipeline.

### Assigning genes to super-enhancers and typical enhancers

We assigned genes to super-enhancers and typical enhancers using a proximity rule as descried in [16,18]. It is known that enhancers tend to loop and communicate with target genes [7], and most of these enhancer-promoter interactions occur within a distance of ~50kb [64]. This approach identified a large proportion of true enhancer/promoter interactions in ESC [65]. Hence, we assigned all transcriptionally active genes to super-enhancers and typical enhancers within a 50kb window.

### Feature-ranking

To find an optimal feature subset, we used two Random Forest based approaches. First, we used Boruta algorithm [37] to rank important features. Briefly, it finds important features by measuring the relevance of each original feature with respect to a reference attribute using Random Forest. Second, we used Random Forest’s out-of-bag approach to calculate the relative importance of each feature. Briefly, this approach takes one feature out and measures its relative importance and contribution to the model. The reason behind to use two approaches to rank features was to see the consistency of feature ranking. We have achieved almost identical results using both methods.

We divided the features into two groups with the aim to achieve two different goals. The 11 transcription factors in mouse embryonic stem cells, were used to rank the transcription factors and explore their importance in super-enhancer prediction. The other 20 chromatin features including, histone modifications (HMs), RNA polymerase II (Pol II), transcriptional co-activating proteins, chromatin regulators (CRs), Cohesin, and sub-units of Lsd1-NuRD complex, were used to ranked these features together in mESC and pro-B cells.

### Motif analysis

We used FIMO (Find Individual Motif Occurrences) version 4.10.0 [66] for motif analysis with p-value < 10^-4^ with a custom library of TRANSFAC motifs including (Oct4: M01124, Sox2: M01272, Nanog: M01123, Esrrb:M01589, Klf4: M01588). The number of occurrences of each motif were counted for the constituents of super-enhancer and typical enhancer regions.

### Gene ontology analysis

We performed the gene ontology analysis using Genomic Regions Enrichment of Annotations Tool (GREAT) web tool (version 3.0.0) [67] with whole-genome as background and default parameters. We reported the top Gene Ontology (GO) terms with the lowest p-value.

### Visualization and statistical analysis

We generated box plots using R programming language by extended the whiskers to 1.5x the interquartile range. The P-values were calculated based on Wilcoxon signed-rank test for box plots, by using wilcox.test function in R. We used ngs.plot (https://github.com/shenlab-sinai/ngsplot) [68], to generate heat maps and normalized binding profiles at the constituents of super-enhancers and typical enhancers and their flanking 3kb regions (for example, Fig. 1a, 4a). We used Intervene tool (http://intervene.readthedocs.io) [69] to generate Venn diagrams.

### Additional files

**Additional file 1**: A PDF document contains Supplementary Figures S1–S4. And also, the public dataset used in this study in Supplementary Table S1-S2.

**Additional file 2**: An Excel spreadsheet contains a list of super-enhancers and typical enhancers and their associated genes in mESC identified by using Cdk8, Cdk9 and Smad3.

**Additional file 3**: An Excel spreadsheet contains a list of super-enhancers and typical enhancers identified by using H3K27ac and Smad3 in pro-B cell.

## Abbreviations

TFs: Transcription Factors
ChIP-seq: Chromatin immune precipitation followed by high-throughput sequencing
ESCs: Embryonic Stem Cells
SEs: Super-enhancers
TEs: Typical enhancers
TSS: Transcription Start Site
SNPs: Single Nucleotide Polymorphisms
GEO: Gene Expression Omnibus
GO: Gene Ontology

## Authors’ contributions

AK conceived and designed the experiments. XG reviewed and approved experiment design. AK performed the experiments and analysed the data. AK wrote the manuscript and XG reviewed it. All authors read and approved the final manuscript.

## Acknowledgements

We thank Anthony Mathelier and Masood ur Rehman Kayani for their comments and suggestions to improve the manuscript. We would like to thank Lydia Danner for proofreading the manuscript. We thank the anonymous reviewers for their useful comments and suggestions.

## References

1. Heintzman ND, Hon GC, Hawkins RD, Kheradpour P, Stark A, Harp LF, et al. Histone modifications at human enhancers reflect global cell-type-specific gene expression. Nature. Nature Publishing Group; 2009;459:108–12.

2. Heinz S, Romanoski CE, Benner C, Glass CK. The selection and function of cell type-specific enhancers. Nat Rev Mol Cell Biol. Nature Publishing Group; 2015;16:144–54.

3. Levine M, Cattoglio C, Tjian R. Looping back to leap forward: transcription enters a new era. Cell. Elsevier Inc.; 2014;157:13–25.

4. Shlyueva D, Stampfel G, Stark A. Transcriptional enhancers: from properties to genome-wide predictions. Nat. Rev. Genet. Nature Publishing Group; 2014;15:272–86.

5. Wamstad J a, Wang X, Demuren OO, Boyer L a. Distal enhancers: new insights into heart development and disease. Trends Cell Biol. Elsevier Ltd; 2014;24:294–302.

6. Kolovos P, Knoch T a, Grosveld FG, Cook PR, Papantonis A. Enhancers and silencers: an integrated and simple model for their function. Epigenetics Chromatin. 2012;5:1.

7. Ong C-T, Corces VG. Enhancer function: new insights into the regulation of tissue-specific gene expression. Nat. Rev. Genet. Nature Publishing Group; 2011;12:283–93.

8. Thurman RE, Rynes E, Humbert R, Vierstra J, Maurano MT, Haugen E, et al. The accessible chromatin landscape of the human genome. Nature. Nature Publishing Group; 2012;489:75–82.

9. Banerji J, Rusconi S, Schaffner W. Expression of a β-globin gene is enhanced by remote SV40 DNA sequences. Cell. 1981;27:299–308.

10. Visel A, Blow MJ, Li Z, Zhang T, Akiyama J a, Holt A, et al. ChIP-seq accurately predicts tissue-specific activity of enhancers. Nature. Nature Publishing Group; 2009;457:854–8.

11. Creyghton MP, Cheng AW, Welstead GG, Kooistra T, Carey BW, Steine EJ, et al. Histone H3K27ac separates active from poised enhancers and predicts developmental state. Proc. Natl. Acad. Sci. U. S. A. 2010;107:21931–6.

12. Natarajan A, Yardimci GG, Sheffield NC, Crawford GE, Ohler U. Predicting cell-type-specific gene expression from regions of open chromatin. Genome Res. 2012;22:1711–22.

13. Kagey MH, Newman JJ, Bilodeau S, Zhan Y, Orlando DA, van Berkum NL, et al. Mediator and cohesin connect gene expression and chromatin architecture. Nature. Nature Publishing Group; 2010;467:430–5.

14. Allen BL, Taatjes DJ. The Mediator complex: a central integrator of transcription. Nat. Rev. Mol. Cell Biol. Nature Publishing Group; 2015;16:155–66.

15. Chen X, Xu H, Yuan P, Fang F, Huss M, Vega VB, et al. Integration of External Signaling Pathways with the Core Transcriptional Network in Embryonic Stem Cells. Cell. 2008;133:1106–17.

16. Whyte WA, Orlando DA, Hnisz D, Abraham BJ, Lin CY, Kagey MH, et al. Master Transcription Factors and Mediator Establish Super-Enhancers at Key Cell Identity Genes. Cell. Elsevier Inc.; 2013;153:307–19.

17. Lovén J, Hoke H a, Lin CY, Lau A, Orlando D a, Vakoc CR, et al. Selective inhibition of tumor oncogenes by disruption of super-enhancers. Cell. 2013;153:320–34.

18. Hnisz D, Abraham BJ, Lee TI, Lau A, Saint-André V, Sigova A a, et al. Super-enhancers in the control of cell identity and disease. Cell. 2013;155:934–47.

19. Pott S, Lieb JD. What are super-enhancers? Nat. Genet. Nature Publishing Group; 2014;47:8–12.

20. Heyn H, Vidal E, Ferreira HJ, Vizoso M, Sayols S, Gomez A, et al. Epigenomic analysis detects aberrant super-enhancer DNA methylation in human cancer. Genome Biol. Genome Biology; 2016;17:11.

21. Lin CY, Erkek S, Tong Y, Yin L, Federation AJ, Zapatka M, et al. Active medulloblastoma enhancers reveal subgroup-specific cellular origins. Nature. Nature Publishing Group; 2016;1–20.

22. Hah N, Benner C, Chong L, Yu RT, Downes M, Evans RM. Inflammation-sensitive super enhancers form domains of coordinately regulated enhancer RNAs. Proc. Natl. Acad. Sci. U. S. A. 2014;

23. Chapuy B, McKeown MR, Lin CY, Monti S, Roemer MGM, Qi J, et al. Discovery and characterization of super-enhancer-associated dependencies in diffuse large B cell lymphoma. Cancer Cell. Elsevier Inc.; 2013;24:777–90.

24. Mansour MR, Abraham BJ, Anders L, Gutierrez A, Durbin AD, Lawton L, et al. An oncogenic super-enhancer formed through somatic mutation of a noncoding intergenic element. Science. 2014;346:1373–7.

25. Ooi WF, Xing M, Xu C, Yao X, Ramlee MK, Lim MC, et al. Epigenomic profiling of primary gastric adenocarcinoma reveals super-enhancer heterogeneity. Nat. Commun. Nature Publishing Group; 2016;7:12983.

26. Parker SCJ, Stitzel ML, Taylor DL, Orozco JM, Erdos MR, Akiyama J a, et al. Chromatin stretch enhancer states drive cell-specific gene regulation and harbor human disease risk variants. Proc. Natl. Acad. Sci. U. S. A. 2013;110:17921–6.

27. Pasquali L, Gaulton KJ, Rodríguez-Seguí S a, Mularoni L, Miguel-Escalada I, Akerman I, et al. Pancreatic islet enhancer clusters enriched in type 2 diabetes risk-associated variants. Nat. Genet. 2014;46:136–43.

28. Vahedi G, Kanno Y, Furumoto Y, Jiang K, Parker SCJ, Erdos MR, et al. Super-enhancers delineate disease-associated regulatory nodes in T cells. Nature. 2015;520:558–62.

29. Witte S, Bradley A, Enright AJ, Muljo SA. High-density P300 enhancers control cell state transitions. BMC Genomics. BMC Genomics; 2015;16:903.

30. Inoue Y, Itoh Y, Abe K, Okamoto T, Daitoku H, Fukamizu a, et al. Smad3 is acetylated by p300/CBP to regulate its transactivation activity. Oncogene. 2007;26:500–8.

31. Pouponnot C, Jayaraman L, Massague J. Physical and Functional Interaction of SMADs and p300/CBP. J. Biol. Chem. 1998;273:22865–9.

32. Wang J, Zhuang J, Iyer S, Lin X, Whitfield TW, Greven MC, et al. Sequence features and chromatin structure around the genomic regions bound by 119 human transcription factors. Genome Res. 2012;22:1798–812.

33. Dekker J. GC- and AT-rich chromatin domains differ in conformation and histone modification status and are differentially modulated by Rpd3p. Genome Biol. 2007;8:R116.

34. Valouev A, Johnson SM, Boyd SD, Smith CL, Fire AZ, Sidow A. Determinants of nucleosome organization in primary human cells. Nature. Nature Publishing Group; 2011;474:516–20.

35. Niederriter A, Varshney A, Parker S, Martin D. Super Enhancers in Cancers, Complex Disease, and Developmental Disorders. Genes. 2015;6:1183–200.

36. Villar D, Berthelot C, Aldridge S, Rayner TF, Lukk M, Pignatelli M, et al. Enhancer Evolution across 20 Mammalian Species. Cell. The Authors; 2015;160:554–66.

37. Kursa MB, Rudnicki WR. Feature Selection with the Boruta Package. J. Stat. Softw. 2010;36.

38. Donner AJ, Ebmeier CC, Taatjes DJ, Espinosa JM. CDK8 is a positive regulator of transcriptional elongation within the serum response network. Nat. Struct. Mol. Biol. Nature Publishing Group; 2010;17:194–201.

39. Feng B, Jiang J, Kraus P, Ng J-H, Heng J-CD, Chan Y-S, et al. Reprogramming of fibroblasts into induced pluripotent stem cells with orphan nuclear receptor Esrrb. Nat. Cell Biol. 2009;11:197–203.

40. Young RA. Control of the embryonic stem cell state. Cell. Elsevier Inc.; 2011;144:940–54.

41. Takahashi K, Yamanaka S. Induction of Pluripotent Stem Cells from Mouse Embryonic and Adult Fibroblast Cultures by Defined Factors. Cell. 2006;126:663–76.

42. Mullen AC, Orlando D a, Newman JJ, Lovén J, Kumar RM, Bilodeau S, et al. Master transcription factors determine cell-type-specific responses to TGF-β signaling. Cell. 2011;147:565–76.

43. Yin J-W, Wang G. The Mediator complex: a master coordinator of transcription and cell lineage development. Dev. Camb. Engl. 2014;141:977–87.

44. Belkina AC, Denis G V. BET domain co-regulators in obesity, inflammation and cancer. Nat. Rev. Cancer. Nature Publishing Group; 2012;12:465–77.

45. Liu W, Ma Q, Wong K, Li W, Ohgi K, Zhang J, et al. Brd4 and JMJD6-associated anti-pause enhancers in regulation of transcriptional pause release. Cell. 2013;155:1581–95.

46. Zhang W, Prakash C, Sum C, Gong Y, Li Y, Kwok JJT, et al. Bromodomain-containing protein 4 (BRD4) regulates RNA polymerase II serine 2 phosphorylation in human CD4+ T cells. J. Biol. Chem. 2012;287:43137–55.

47. Itzen F, Greifenberg AK, Bösken C a., Geyer M. Brd4 activates P-TEFb for RNA polymerase II CTD phosphorylation. Nucleic Acids Res. 2014;42:7577–90.

48. Di Micco R, Fontanals-Cirera B, Low V, Ntziachristos P, Yuen SK, Lovell CD, et al. Control of Embryonic Stem Cell Identity by BRD4-Dependent Transcriptional Elongation of Super-Enhancer-Associated Pluripotency Genes. Cell Rep. 2014;1–14.

49. Pelish HE, Liau BB, Nitulescu II, Tangpeerachaikul A, Poss ZC, Silva DH Da, et al. Mediator kinase inhibition further activates super-enhancer-associated genes in AML. Nature. 2015;526:273–276.

50. Hnisz D, Schuijers J, Lin CY, Weintraub AS, Abraham BJ, Lee TI, et al. Convergence of Developmental and Oncogenic Signaling Pathways at Transcriptional Super-Enhancers. Mol. Cell. Elsevier Inc.; 2015;1–9.

51. Carey M. The enhanceosome and transcriptional synergy. Cell. 1998;92:5–8.

52. Corradin O, Saiakhova A, Akhtar-Zaidi B, Myeroff L, Willis J, Cowper-Sallari R, et al. Combinatorial effects of multiple enhancer variants in linkage disequilibrium dictate levels of gene expression to confer susceptibility to common traits. Genome Res. 2014;24:1–13.

53. Dowen JM, Fan ZP, Hnisz D, Ren G, Abraham BJ, Zhang LN, et al. Control of Cell Identity Genes Occurs in Insulated Neighborhoods in Mammalian Chromosomes. Cell. Elsevier Inc.; 2014;159:374–87.

54. Siersbæk R, Rabiee A, Nielsen R, Sidoli S, Traynor S, Loft A, et al. Transcription factor cooperativity in early adipogenic hotspots and super-enhancers. Cell Rep. The Authors; 2014;7:1443–55.

55. Hay D, Hughes JR, Babbs C, Davies JOJ, Graham BJ, Hanssen LLP, et al. Genetic dissection of the α-globin super-enhancer in vivo. Nat. Genet. Nature Publishing Group; 2016;1–12.

56. Shin HY, Willi M, Yoo KH, Zeng X, Wang C, Metser G, et al. Hierarchy within the mammary STAT5-driven Wap super-enhancer. Nat. Genet. 2016;48:904–11.

57. Dukler N, Gulko B, Huang Y, Siepel A. Is a super-enhancer greater than the sum of its parts? 2017;49:2–7.

58. Ouyang Z, Zhou Q, Wong WH. ChIP-Seq of transcription factors predicts absolute and differential gene expression in embryonic stem cells. Proc. Natl. Acad. Sci. U. S. A. 2009;106:21521–6.

59. Khan A, Zhang X. dbSUPER: a database of super-enhancers in mouse and human genome. Nucleic Acids Res. 2016;44:D164–D171.

60. Rosenbloom KR, Armstrong J, Barber GP, Casper J, Clawson H, Diekhans M, et al. The UCSC Genome Browser database: 2015 update. Nucleic Acids Res. 2014;43:D670–81.

61. Langmead B, Trapnell C, Pop M, Salzberg SL. Ultrafast and memory-efficient alignment of short DNA sequences to the human genome. Genome Biol. 2009;10:R25.

62. Rahl PB, Lin CY, Seila AC, Flynn R a., McCuine S, Burge CB, et al. C-Myc regulates transcriptional pause release. Cell. 2010;141:432–45.

63. Zhang Y, Liu T, Meyer C a, Eeckhoute J, Johnson DS, Bernstein BE, et al. Model-based analysis of ChIP-Seq (MACS). Genome Biol. 2008;9:R137.

64. Chepelev I, Wei G, Wangsa D, Tang Q, Zhao K. Characterization of genome-wide enhancer-promoter interactions reveals co-expression of interacting genes and modes of higher order chromatin organization. Cell Res. Nature Publishing Group; 2012;22:490–503.

65. Dixon JR, Selvaraj S, Yue F, Kim A, Li Y, Shen Y, et al. Topological domains in mammalian genomes identified by analysis of chromatin interactions. Nature. Nature Publishing Group; 2012;485:376–80.

66. Grant CE, Bailey TL, Noble WS. FIMO: scanning for occurrences of a given motif. Bioinforma. Oxf. Engl. 2011;27:1017–8.

67. McLean CY, Bristor D, Hiller M, Clarke SL, Schaar BT, Lowe CB, et al. GREAT improves functional interpretation of cis-regulatory regions. Nat. Biotechnol. Nature Publishing Group; 2010;28:495–501.

68. Shen L, Shao N, Liu X, Nestler E. ngs.plot: Quick mining and visualization of next-generation sequencing data by integrating genomic databases. BMC Genomics. BMC Genomics; 2014;15:284.

69. Khan A, Mathelier A. Intervene: a tool for intersection and visualization of multiple gene or genomic region sets. BMC Bioinformatics. 2017;18:287.

